# Hippocampal Projections to the Striatal Compartments, Striosome and Matrix, are Spatially Segregated in CA1

**DOI:** 10.64898/2026.01.27.702163

**Authors:** An N. Tieu, Alishba Sadiq, Jeff L. Waugh

## Abstract

The hippocampus routes information to the striatum through at least four polysynaptic circuits. Striatal projection neurons are organized into two tissue compartments, the matrix and striosome, which differ in their embryologic origins, relative abundance, intra-striate location, and afferent and efferent connectivity. These compartments are embedded in distinct functional networks and are activated by different tasks. Consequently, hippocampal inputs that route preferentially through the striosome may underpin different functions and engage with different remote networks than inputs that route through the matrix. It was unknown whether striosome-bound and matrix-bound projections from the hippocampus followed different polysynaptic circuits. We assessed hippocampo-striate projections in living humans using probabilistic diffusion tractography by first parcellating the striatum into voxels with striosome-like and matrix-like structural connectivity. We then quantified structural connectivity between hippocampal efferents (CA1) to each set of compartment-like voxels. CA1 projections to striosome-like voxels in the dorsal striatum (caudate and putamen) were 3.1-fold more abundant than those to matrix-like voxels, particularly in caudo-lateral CA1. This striosome-favoring bias was similar in three segregated hippocampo-striate circuits, in streamlines routing through the subiculum, lateral septum, or medial prefrontal cortex. However, a small region in rostro-medial CA1 preferentially targeted matrix-like voxels. Functional connectivity between CA1 and compartment-like voxels matched this segregated pattern: CA1 activation was correlated with striosome-like voxels but anti-correlated with matrix-like voxels. Additionally, streamlines from CA1 to nucleus accumbens exhibited hemispheric asymmetries, with the left hemisphere biased towards matrix and the right towards striosome. These findings suggest that hippocampo-striate projections are spatially segregated into compartment-specific circuits.

## Introduction

The hippocampus and striatum have complementary and interactive roles in memory and learning (Mizumori, Yeshenko et al. 2004, Amso, Davidson et al. 2005, Rossato, Zinn et al. 2006). The hippocampus and its CA1 subfield are integral to declarative memory encoding and retrieval, supporting episodic and associative memories of past events and facts (Morris, Garrud et al. 1982, Schlichting and Preston 2014). In contrast, the striatum is often linked to habitual memories, characterized by stimulus-response associations and behavior reinforcement (Brasted, Humby et al. 1997, Poldrack and Rodriguez 2004). These systems can function independently or in parallel, operating either cooperatively (Kim and Baxter 2001) or competitively (Poldrack and Rodriguez 2004) to guide behavior based on prior experience (Packard, Kohlmaier et al. 1996). Selective inhibition of the hippocampus or striatum in animals demonstrates that maze tasks can be completed using either system alone or in combination (Ghiglieri, Gambarana et al. 1997). These manipulations indicate that initial learning is primarily hippocampus-dependent, whereas the transition to a well-learned, habitual behavior involves the striatum (Ghiglieri, Gambarana et al. 1997). Integrating these two systems through effective cross-communication is thought to be essential for normal memory performance (Voermans, Petersson et al. 2004, Müller, Konrad et al. 2018). Delineating the neural circuitry that mediates information transfer from the hippocampus to both the dorsal (caudate and putamen) and ventral (nucleus accumbens) striatum is therefore critical for understanding how these regions support learning and memory. Although the projections linking hippocampus to striatum have been described in animals, it is unknown whether human hippocampo-striate connectivity follows these patterns, and how these projections might be segregated to distinct tissue compartments within the striatum.

Projection neurons in the striatum are organized into two distinct tissue compartments, the matrix and striosome. The striosome is a web-like structure that is interdigitated with and surrounded by the matrix (Brimblecombe and Cragg 2016). These compartments differ in the expression of histochemical markers (Crittenden, Tillberg et al. 2016), their spatial distribution within the striatum (Graybiel and Ragsdale 1978), and their relative abundance (Graybiel and Ragsdale 1978, Holt, Graybiel et al. 1997). Matrix and striosome are embedded in segregated structural (Funk, Hassan et al. 2023, Funk, Hassan et al. 2024) and functional networks (Sadiq, Funk et al. 2025). This suggests that information routed through one compartment versus the other may regulate discrete brain functions. Each compartment is also activated or inactivated at different task phases to facilitate specific behavioral and learning outcomes. For example, in mice, striosomal neuron activity was increased during reward and punishment learning (Bloem, Huda et al. 2017). Activation of striosomal neurons via chemogenetic stimulation in mice decreases contralateral turning and distance moved (Okunomiya, Watanabe et al. 2025). In rats, electrical stimulation of the striosome, but not the matrix, induces addiction-like behaviors, such as repetitive lever pressing (White and Hiroi 1998). Evidence that the matrix and striosome are differentially involved in behavior and learning makes it critical to determine whether hippocampo-striate projections exhibit compartment-specific biases. Characterizing this structural connectivity will establish a mechanistic framework for understanding hippocampus-striatum interactions in both healthy cognition and disease. For example, in early Huntington disease atrophy is relatively selective for the striosome compartment (Tippett, Waldvogel et al. 2007), a disease phase in which multiple types of memory are impaired (Zhang, Shen et al. 2022). Mapping compartment-specific hippocampal–striatal pathways could therefore reveal how disruptions in these circuits contribute to disease-related symptoms.

We developed a method for identifying compartment-like voxels in the striatum via connectivity-based parcellation using probabilistic diffusion tractography (Waugh, Hassan et al. 2022). Injected tract tracer studies in animals (reviewed in Waugh, 2022), and our prior neuroimaging-based tract mapping in humans (Funk, Hassan et al. 2023, Funk, Hassan et al. 2024) demonstrated that many extra-striate regions are biased toward either matrix or striosome. Differential connectivity to these compartment-biased extra-striate regions can identify voxels with matrix-like or striosome-like structural connectivity. Voxels identified with this method recapitulate the anatomic features of matrix and striosome demonstrated in animal and human tissue: their relative abundance (Holt, Graybiel et al. 1997), their somatotopic organization (Flaherty and Graybiel 1993), and their spatial distribution (Graybiel and Ragsdale 1978). Connectivity-based parcellation selects a precise set of striatal voxels; shifting voxel location by 2-3 voxels at random eliminates compartment-specific bias (Funk, Hassan et al. 2024, Sadiq, Funk et al. 2025). In subjects scanned twice, parcellated striatal voxels had a low test– retest error rate (0.14%) (Waugh, Hassan et al. 2022).

Viral tract tracing in mice revealed that polysynaptic pathways link CA1 to the striatum (Du, Li et al. 2023). Du et al. demonstrated that CA1 connects to both the dorsal and ventral striatum via intermediate nuclei. The subiculum serves as a major relay linking dorsal CA1 to the dorsal and ventral striatum, whereas the entorhinal and medial prefrontal cortices mediate ventral CA1 projections to the same targets. The lateral septum also provides a connection from dorsal CA1 to the nucleus accumbens and from ventral CA1 to the dorsal striatum. In contrast, viral tracers demonstrated that retrosplenial cortex and supramammillary nucleus receive prominent projections from CA1 but do not relay projections to either dorsal or ventral striatum. While Du et al.’s viral-tracing work mapped these pathways in mice, characterization of these circuits in living humans requires diffusion-based inference rather than direct anatomical tracing. We set out to characterize these polysynaptic pathways in living humans, using probabilistic diffusion tractography to quantify structural connectivity from CA1 to matrix-like and striosome-like voxels via the intermediary nuclei identified by Du et al. Quantifying structural connectivity from CA1 to compartment-like voxels allowed us to determine whether hippocampo-striate projections exhibit compartmental biases, and how those projections were arrayed among the polysynaptic circuits of Du et al. We then assessed the functional significance of compartment-selective biases in structural connectivity by correlating resting-state functional activation in CA1 with activation in matrix-like and striosome-like voxels.

We found that in humans, structural connectivity in the three CA1-dorsal striatum pathways (subicular, medial prefrontal, and lateral septal paths) identified by Du et al. (Du, Li et al. 2023) was markedly biased toward striosome-like voxels. Voxelwise analyses revealed that this striosome-like bias was localized to the lateral and caudal regions of CA1, whereas a smaller, distinct region in the medial- and rostral-most CA1 was biased toward matrix-like voxels. Resting state functional connectivity networks between CA1 and the compartments (separate MRI scans in the same subjects) identified a similar pattern: activity in CA1 was positively correlated with striosome-like voxels and anti-correlated with matrix-like voxels. These results provide the first *in-vivo* evidence that hippocampal projections to the human striatum exhibit compartment-level segregation, and that compartment-selective bias in structural connectivity may underpin opposing hippocampal networks for striosome and matrix. Segregating hippocampo-striate projections into striosome- or matrix-biased networks may be a structural basis for the competing and complementary functions of the hippocampus and striatum.

## Methods

### Overview

We first parcellated the striatum into matrix-like and striosome-like masks via a connectivity based parcellation. We then performed subsequent rounds of tractography (traditional streamline and classification targets) to assess connectivity between CA1 and the parcellated compartment-like masks. We also assessed resting state functional connectivity between the compartment-like masks and compartment-specific regions within CA1.

### MRI Acquisition

This was a secondary analysis of diffusion MRI data obtained from the Human Connectome Project (HCP) S1200 data release (Van Essen, Smith et al. 2013). All participants provided written consent to participate in the original HCP protocols, and to the sharing of their data for secondary analyses.

All subjects were scanned on Siemens 3T scanners with imaging protocols and acquisition parameters standardized across multiple sites (Van Essen, Smith et al. 2013). Diffusion tensor imaging (DTI) scans were acquired at a resolution of 1.25 mm isotropic using 200 directions (14 B0 volumes, 186 volumes at noncolinear directions) and the following parameters: repetition time = 3.23 s, echo time = 0.0892. DTI scans included both anterior-posterior and posterior-anterior acquisitions, allowing for correction of susceptibility artifacts. T1 weighted scans were collected at a 0.80 × 0.76 × 0.76 mm resolution. Complete MRI scan protocols can be found in the HCP S1200 Subject Data Release Reference Manual.

Resting-state fMRI was acquired in four runs, with two runs collected per session across two separate sessions. Within each session, phase-encoding direction alternated between right-to-left (RL) and left-to-right (LR) acquisitions. During scanning, participants were instructed to keep their eyes open and fixated on a central crosshair. For the present study, we analyzed data from the first resting-state session, concatenating the RL and LR runs to yield a total of 2,400 timepoints (1,200 per run). Acquisition parameters were as follows: TR = 720 ms, TE = 33 ms, flip angle = 52°, 72 slices, 2.0 mm isotropic resolution, with a total scan duration of 14 minutes and 33 seconds per run.

### MRI Preprocessing

We processed DTI data using the FSL FMRIB Diffusion Toolbox with standard parameters (Smith, Jenkinson et al. 2004). We performed skull stripping and brain extraction using *bet2* and corrected for eddy current distortion and subject movement during imaging acquisition using *eddy_cuda10*.*2* (Andersson and Sotiropoulos 2016). We used *dtifit* to assign diffusion tensors to each voxel and *bedpostx_gpu* to model crossing fibers and calculate diffusion probability estimates at each voxel (Hernández, Guerrero et al. 2013). We generated registration matrices from the FMRIB58_FA standard diffusion space to each subject’s native diffusion space using *flirt* and *fnirt*. We used these matrices to register all MNI-space regional segmentations into each subject’s native diffusion space.

We segmented all regions of interest used for striatal parcellation based on the MNI152_T1_1 mm standard brain using the atlases of Talairach and Tournoux (Talairach and Tournoux 1988). We used the FreeSurfer automated segmentation tool *recon-all*, including its hippocampal subfields module, to segment the hippocampal CA1 region in each subject’s native T1 space (Iglesias, Augustinack et al. 2015). The dorsal striatum mask included the caudate and putamen but excluded the posterior half of the caudate tail, as this region contains very little striosome (Bernácer, Prensa et al. 2008) and its narrow structure in the coronal plane reduced registration accuracy due to partial volume effects with the surrounding white matter.

### Identifying Matrix-like and Striosome-like Voxels

The striosome is uniquely situated in each individual, making region-of-interest masks inaccurate for assessing the striatal compartments. Instead, we used probabilistic diffusion tractography to identify subject-specific compartment-like masks via connectivity-based parcellation (Waugh, Hassan et al. 2022). Matrix and striosome tissue have structural connectivity biases with extra-striate regions, previously identified through decades of tract-tracing studies in animals and imaging in humans (reviewed in Waugh, 2022). We identified voxels as matrix-like or striosome-like based on their structural connectivity patterns to these extra-striate bait regions. We first created matrix- and striosome-favoring bait region masks. The matrix-favoring mask consisted of the supplementary motor area, primary motor cortex, primary sensory cortex (Brodmann areas 1-3), the combined ventrolateral thalamic nuclei, and the globus pallidus interna (Malach and Graybiel 1986, Jimenez-Castellanos and Graybiel 1989, Gimenez-Amaya and Graybiel 1990, Ragsdale and Graybiel 1991, Ebrahimi, Pochet et al. 1992, Parthasarathy, Schall et al. 1992, Flaherty and Graybiel 1993, Desban, Gauchy et al. 1995, Inase, Sakai et al. 1996, Kincaid and Wilson 1996, McGregor, McKinsey et al. 2019). The striosome-favoring mask consisted of the posterior orbitofrontal cortex, basal operculum, anterior insula, basolateral amygdala, and mediodorsal thalamus (Berendse, Voorn et al. 1988, Ragsdale and Graybiel 1990, Ragsdale and Graybiel 1991, Eblen and Graybiel 1995, Unzai, Kuramoto et al. 2015). These bait regions matched those we utilized with other human MRI datasets (Waugh, Hassan et al. 2022, Funk, Hassan et al. 2024, Waugh, Hassan et al. 2025). Readers should note that while these structural connectivity biases are well established for the dorsal striatum, we could identify no injected tracer studies that assessed connectivity with the nucleus accumbens. In the absence of any histologic findings to guide the selection of accumbens-specific bait regions, we utilized the same compartment-favoring masks in dorsal and ventral striatal parcellations.

We performed tractography using FSL *probtrackx2_gpu* (Hernandez-Fernandez, Reguly et al. 2019) in classification targets mode with the following parameters: curvature threshold=0.2; steplength=0.5 mm; 2,000 steps per sample; 5,000 streamlines per seed voxel, setting striatal voxels as the seed and the matrix- or striosome-favoring bait regions as targets. All rounds of tractography were performed in native diffusion space, and separately in each hemisphere. We classified each striatal voxel as matrix-like, striosome-like, or indeterminant depending on the ratio of streamlines to the matrix- or striosome-favoring bait masks.

Probabilistic tractography is influenced by the volume of the masks utilized as targets for streamlines. We generated equal-volume matrix-like and striosome-like masks to avoid this source of bias. We identified the voxels with the largest compartment-specific bias in the matrix-like and striosome-like probability distributions, separately for the dorsal striatum (caudate and putamen) and nucleus accumbens. We set the target size for these compartment-like masks equal to the volume of either striatum or nucleus accumbens at 1.5 standard deviations above the mean (13% of the volume of that nucleus). For cases in which a subject’s compartment-like volume was smaller than the target volume, we set the volume target of both compartments to that of the smaller mask; the volume of the matrix-like and striosome-like masks was always equal. To minimize sampling variability in the (smaller) nucleus accumbens, subjects were excluded from nucleus accumbens analyses if compartment volumes were below 5 mm^3^. After removing subjects with inadequate mask sizes, the subject count for analyses involving the nucleus accumbens was 546 subjects. We describe compartment voxels as “-like” because striatal parcellation identifies voxels whose structural connectivity patterns match the connectivity biases of matrix and striosome that were identified in tissue. However, striatal parcellation should not be conflated with direct identification of the compartments through immunohistochemical staining, the gold standard for identifying matrix and striosome.

### Tractography Experiments

After we parcellated each individual’s striatum, we performed 10 distinct rounds of tractography to examine the connectivity between CA1 with the dorsal striatum or nucleus accumbens (Table 1). We used *probtrackx2_gpu* in both classification targets tractography (CTT) mode and traditional streamline tractography. When performing CTT, we used the same parameters described above for striatal parcellation. For streamline tractography, we generated a denser probability function by increasing the streamlines per seed voxel by 10-fold, to 50,000. This reduced the intra-subject variance in streamline amplitude. We performed tractography separately for each hemisphere by using a block mask to prevent inter-hemispheric streamlines. For all rounds of tractography we excluded streamlines that propagated through the supramammillary nucleus, retrosplenial cortex, or amygdala, regions that were not intermediate relays for polysynaptic connections from CA1 to striatum (Du, Li et al. 2023). Excluding these regions reduced low-fidelity streamlines, improving anatomical precision.

**Table 1:** We carried out 10 rounds of probabilistic tractography to address distinct experimental goals. CTT: Classification Targets tractography, NAcc: Nucleus Accumbens, LS: Lateral Septum, mPFC: medial Prefrontal Cortex, SUB: Subiculum.

| Round | Type of Tractography | Seed | Targets | Goal of Tractography |
| --- | --- | --- | --- | --- |
| 1 | CTT | CA1 | Caudate or Putamen | Assess for caudate or putamen biases in connectivity from CA1 |
| 2 | CTT | CA1 | Matrix-like or Striosome-like voxels | Assess for compartment-like biases in connectivity from CA1 |
| 3 | CTT | CA1 | Matrix-near-Striosome or Striosome-like masks | Control: tests if the bias towards striosome-like voxels was a somatotopic effect |
| 5 | CTT | CA1 | Compartment-like masks in Striatum with either the LS, mPFC, or SUB as obligatory waypoints | Assess for compartment-like biases in connectivity from CA1 via the previously identified relay regions |
| 6 | CTT | Striatum | CA1 | Quantify the Dice similarity coefficient of striatal voxels that route through SUB vs mPFC |
| 7 | Streamline | Striatum | Matrix-favoring or Striosome-favoring regions of CA1 | Assess the compartment-bias of striatal voxels with high-connectivity to CA1 |
| 8 | CTT | CA1 | Matrix-like or Striosome-like voxels | Assess for compartment-like biases in connectivity from CA1 |
| 9 | CTT | CA1 | Compartment-like masks in Nucleus Accumbens with either the LS, mPFC, or SUB as obligatory waypoints | Assess for compartment-like biases in connectivity from CA1 via the previously identified relay regions |
| 10 | Streamline | CA1 | Matrix-like or Striosome-like voxels in dorsal striatum/nucleus accumbens | Assess the degree to which compartment bias depends on the most dominant route |

#### 1. CA1 to Dorsal Striatum Nucleus Bias

In round one, we assessed connectivity between CA1 and the caudate vs. putamen using classification targets tractography, independent of a bias toward matrix- or striosome-like voxels. Given that the caudate and putamen have different volume and the distributions of matrix and striosome differ between them (Graybiel and Ragsdale 1978, Brimblecombe and Cragg 2016), biases in connectivity between CA1 and the nuclei could skew assessments of the matrix and striosome within them. We identified such a nuclear bias, leading us to generate compartment-like masks such that their volumes were proportional to the relative volumes of caudate and putamen, reducing the influence of this CA1-nucleus bias.

#### 2. CA1 to Compartment Bias: Dorsal Striatum

In round two, we assessed for biases in connectivity between CA and the proportionate compartment-like masks in the caudate and putamen using classification targets tractography. We seeded streamlines from CA1 voxels, with the matrix-like and striosome-like masks as targets.

#### 3. Assessing for a Neighborhood Effect in CA1 to Dorsal Striatum Compartment Bias

In the third round of tractography, we asked whether CA1-compartment biases from round two were specific for matrix-like and striosome-like masks, or if these results instead reflected differences in CA1’s connectivity with different regions of the striatum. Since striosome and matrix are enriched in different parts of the striatum, it is appropriate to question whether differences in structural connectivity are mediated by compartment, or by regional differences in connectivity within the striatum, independent of compartment. For example, if CA1-striatum projections were somatotopically biased (e.g., toward rostro-ventral striatum, where striosome is enriched), this could bias measures of CA1-compartment connectivity. Therefore, we designed this control experiment to learn whether CA1 was biased toward a striatal “neighborhood” or toward our precisely-selected compartment-like voxels. We created new matrix masks (matrix-near-striosome) by identifying highly biased matrix-like voxels that were restricted to a 5 mm radius surrounding the previously-defined striosome-like mask.

That is, we identified the most-biased matrix-like voxels within the neighborhood of a striosome-like voxel instead of identifying the most-biased matrix-like voxels anywhere in the striatum. We began this search within a 3 mm radius from a striosome-like voxel and expanded to 4 mm, then 5 mm only if a subject did not have adequate matrix-like volume to reach our target voxel volume at more restrictive thresholds. We did not resample striosome-like voxels because their mask volume (13% of the striatum) used up nearly all of the expected volume of striosome tissue (15% of the striatum) – there were too few unused striosome-like voxels to pick masks that differed from the original striosome-like masks. We performed classification targets tractography targeting the matrix-near-striosome vs. the original striosome masks.

#### 4. CA1 to Dorsal Striatum, Route-Specific Compartment Bias

In our fifth round of tractography, we sought to replicate the pathways observed by Du et al. (Du, Li et al. 2023), by adding either the subiculum (SUB), lateral septum (LS), or medial prefrontal cortex (mPFC) as an obligatory waypoint for streamline generation. When testing one intermediary waypoint, we excluded the other two intermediary waypoints to select for streamlines that followed that specific path. We did not assess the entorhinal cortex, the fourth of Du et al.’s relay regions, because streamline tractography cannot distinguish fibers originating in the subiculum, but passing through the entorhinal cortex, from those originating in the entorhinal cortex.

#### 5. CA1 to Dorsal Striatum, Route-Specific Dice Similarity Coefficient

In the sixth round of tractography, we assessed the somatotopy of CA1-striatum projections that routed through the SUB and mPFC routes. We did not include the LS route due to its similarity with the SUB route in preceding comparisons. We assessed overlap within each individual’s striatum, in native diffusion space, using the Dice similarity coefficient (DSC) to compare striatal streamline density maps for the SUB and mPFC routes. We performed CTT, seeding from all striatal voxels and targeting CA1, and including either SUB or mPFC as a waypoint and excluding the other region. We separated each map into high- and low-connectivity voxels – the uppermost 25^th^ percentile of streamline counts vs. the lowest 75^th^ percentile – using *fslmaths* (-thrP 75). We calculated the DSC for the two high-connectivity masks (SUB vs. mPFC), and for the two low-connectivity masks (SUB vs. mPFC). We also extracted the mean compartment bias within the high- and low-connectivity voxels for both routes.

#### 6. CA1 to Dorsal Striatum, Verification of Striosome-bias with High-Connectivity Voxels

In the sixth tractography round, we examined whether voxels exhibiting high connectivity between CA1 and the striatum preferentially routed to matrix-favoring versus striosome-favoring subregions of CA1 along the SUB and mPFC pathways. We also assessed whether streamlines projecting to the matrix-favoring and striosome-favoring regions of CA1 followed distinct routes. We performed streamline tractography, seeding from the whole-striatum and targeting the matrix-favoring and striosome-favoring parts of CA1 (established in tractography round 2). We thresholded the streamline density maps to extract high connectivity voxels or voxels with large streamline counts (uppermost 25^th^ percentile streamline counts) and measured the overlap of voxels which routed to either the matrix-favoring or striosome-favoring zones.

#### 7. CA1 to Nucleus Accumbens Compartment Bias

We assessed connectivity between CA1 and compartment-like voxels in the nucleus accumbens using CTT. We seeded CA1 and targeted the matrix-like and striosome-like voxels in the nucleus accumbens. Round 8 was a direct correlate of Round 2; parameters were identical except for the switch from striatum to nucleus accumbens targets.

#### 8. CA1 to Nucleus Accumbens, Route-Specific Compartment Bias

This round of tractography replicated in the nucleus accumbens what we investigated in the dorsal striatum in round 5. We added the previously-described relay nuclei (SUB, LS, mPFC) as obligatory waypoints to assess the circuitry described by Du et al. (Du, Li et al. 2023). When performing tractography using an intermediary waypoint, the other two intermediary regions were added as exclusion masks.

#### 9. Assessing the Degree to which Compartment-like Bias Depends on the Dominant Route of Streamlines

We examined the degree to which the observed striosome-bias in connectivity between CA1 and the dorsal striatum/nucleus accumbens depended on the dominant streamline bundle. We performed streamline tractography, seeding from CA1 and targeting the compartment-like masks. We averaged subject-level streamline density maps and used a percentile-based threshold to define tract regions contributing most strongly to CA1-dorsal striatum/nucleus accumbens connectivity. Probabilistic tractography produces streamlines that contact virtually every voxel in the brain, with the majority including only a negligible number of streamlines. These lower amplitude voxels are likely to be a result of contacts from extraneous non-anatomic streamlines. However, because the streamlines at these non-specific voxels were sparse compared to voxels near the center of the tract, we were able to use thresholding at the individual level to exclude these non-specific voxels from our analyses (Waugh, Hassan et al. 2022). We applied an amplitude threshold to include voxels within the top 95% of streamline counts, thereby segmenting the dominant tract. We added this region as an exclusion mask for CTT, seeding from CA1 and targeting the matrix-like and striosome-like masks.

#### Compartment-Specific CA1 Functional Connectivity Analysis

We used matrix-like and striosome-like striatal masks, defined through structural connectivity-based parcellation (see Section on Identifying Matrix-like and Striosome-like Voxels), as seeds for resting state functional connectivity with CA1. In each hemisphere, we extracted resting-state fMRI time series from the CA1 ROIs and from striosome-like and matrix-like masks. We computed Pearson correlations between the BOLD time series for functional activation in CA1 and compartment-like striatal masks, then Fisher z-transformed these correlations. We calculated CA1 functional connectivity values separately for striosome-like and matrix-like masks in each hemisphere. These values were then averaged across subjects to generate group-level estimates of compartment-specific CA1 connectivity. We used paired-sample *t*-tests to directly compare CA1 functional connectivity between striosome-like and matrix-like masks within each hemisphere, allowing assessment of both the direction and magnitude of compartment-specific coupling.

#### Statistical Analyses

We quantified compartment-like biases in connectivity by extracting the volume of CA1 whose probability of connection (the ratio of streamlines from “seed to target” for each compartment-like distribution) was ≥ 0.55. We performed two-tailed, two-sample t-tests with unequal variance to assess for statistical significance in our compartment bias measures and DSC’s. In addition to these t-tests, we used the FSL tool *randomise* to perform voxel-wise testing of the CA1 mask to identify voxel clusters with significant differences in compartment-like bias. We performed *randomise* with 5,000 permutations, 2 mm variance smoothing, threshold-free cluster enhancement and combined the left and right hemispheres. Age and sex were included as covariates within *randomise* and regressed as nuisance variables.

## Results

### Experimental Cohort

This study included 921 healthy young adults, comprising 509 females and 412 males, with an average age of 28.7 years (range: 22.0-37.0). Right-handed subjects numbered 843, while 78 were left-handed.

### Structural Connectivity in CA1 is Biased Toward Caudate Over Putamen

We performed CTT, seeding from CA1 and targeting the caudate or putamen (without selecting for compartment-like voxels). The volume of CA1 routing to the caudate was 2.9-fold greater than the volume routing to putamen (362 vs. 124 mm^3^, respectively; p<10^-100^). Differences in the abundance of matrix and striosome within the nuclei (Graybiel and Ragsdale 1978, Holt, Graybiel et al. 1997), coupled with this CA1 bias toward caudate over putamen, could influence further assessments of compartment-specific bias. To control for this CA1-caudate bias, we generated compartment-like masks that were proportionate to the relative volume of caudate or putamen, matching the volume of matrix-like and striosome-like voxels within caudate and within putamen. All subsequent assessments of bias in CA1-dorsal striatum connectivity used these proportionate compartment-like masks.

### Structural Connectivity for CA1 to Dorsal Striatum is Biased Towards Striosome-like Voxels

We performed CTT, seeding from CA1 and targeting compartment-like voxels in caudate and putamen. CA1 voxels that were biased toward striosome-like masks were 3.1-fold more abundant than those biased toward matrix-like voxels (striosome: 386 mm^3^ vs. matrix: 125 mm^3^; p<10^-100^). Both the left (striosome: 372 mm^3^, matrix: 125 mm^3^; p<10^-100^) and right (striosome: 398 mm^3^, matrix: 125 mm^3^; p<10^-100^) hemispheres exhibited the same pattern of increased striosome-like bias. A total volume of 53.5 mm^3^ fell within the indeterminate compartment bias range (0.45 < P < 0.55). Voxelwise assessment of bias (using FSL’s *randomise*, Figure 1) identified a symmetrical segment of caudo-lateral CA1 that was significantly biased toward striosome-like voxels (mean probability of connection to striosome-like masks: 0.70), whereas a smaller rostro-medial cluster was biased toward matrix-like voxels (mean probability of connection to matrix-like masks: 0.59; striosome-favoring vs. matrix-favoring volume, p=7.9×10^-92^).

**Figure 1:**
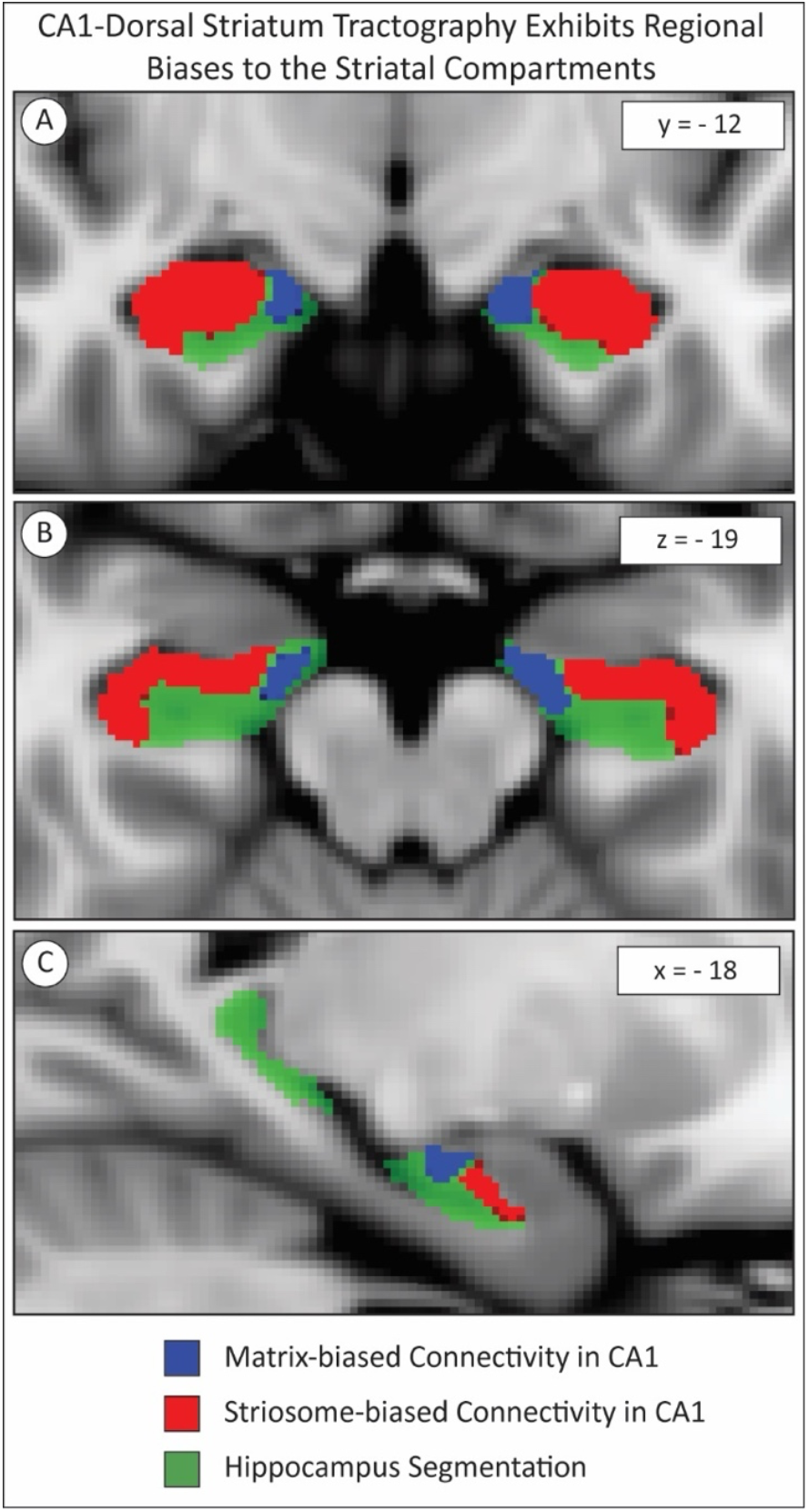
CA1 subregions were biased toward matrix-like (blue) or striosome-like (red) voxels in the dorsal striatum (caudate and putamen), as shown in the coronal (A), axial (B), and sagittal (C) views. Tractography revealed that the inferior-lateral portion of CA1 preferentially projects to striosome-like voxels, whereas a smaller rostral-medial portion of CA1 preferentially projects to matrix-like voxels. Significance threshold, p< 0.05, FWE-corrected within *randomise*. The images adhere to radiographic convention and coordinates follow MNI convention.

### CA1 to Striatum Striosome bias is Dependent on the Core Streamline Bundle

We examined the degree to which the striosome-bias in connectivity between CA1 and dorsal striatum was the result of specific vs. non-specific streamlines. We excluded streamlines that passed through the dominant streamline bundle to select for non-specific, anatomically-implausible streamlines. This led to a 7.9-fold increase in the volume of indeterminate voxels and a reversal in compartment-specific bias in CA1: the mean volume of CA1 biased towards matrix-like masks was 113 mm^3^ and toward striosome-like masks was 29.7 mm^3^ (p<10^-100^). Indirect and implausible streamlines, outside of the core streamline bundle linking CA1 to dorsal striatum, resulted in the elimination of CA1’s striosome-favoring bias.

### The Striosome-Connectivity Bias is not a Neighborhood Effect

Matrix and striosome are enriched in different parts of the striatum (Holt, Graybiel et al. 1997). The compartment-specific bias in CA1-striatal connectivity we described could result from increased connectivity with striosome-like voxels, or it could arise from differences in the strength of connectivity with different regions of the striatum. For example, striosome is enriched in the rostral, medial, and ventral striatum. If CA1-striatum connectivity was enriched to those same regions of the striatum, independent of compartment, this could falsely suggest a bias toward striosome-like voxels. To test whether the apparent striosome bias reflected streamlines targeting the “neighborhood” of striosome-like voxel, rather than the striosome-like voxels themselves, we performed CTT from CA1 to two targets: the original striosome-like masks and a newly generated “matrix-near-striosome” mask (mean matrix bias: P= 0.79). The matrix-near-striosome mask replaces the original matrix-like mask (mean matrix bias: P=0.97). The matrix-near-striosome mask consisted of matrix-like voxels in a zone immediately surrounding striosome-like voxels, in contrast with the original matrix-like voxels that could be located in any part of the striatum. The volume of CA1 with bias toward matrix-near-striosome voxels was 127 mm^3^, whereas 349 mm^3^ routed to striosome-like voxels (p<10^-100^). A volume of 86.1 mm^3^ was within the indeterminate compartment bias range (0.45 < P < 0.55). The striosome-favoring bias from this tractography was not meaningfully different from our original bias assessment, in which matrix-like voxels could reside in any striatal “neighborhood.” This confirms that striosome-biased structural connectivity in CA1 is specific to the precisely selected compartment-like mask, not to the striatal “neighborhood” in which they reside.

### Route-specific Biases in Structural Connectivity: CA1 to Striatal Compartment-like Voxels

To investigate the polysynaptic hippocampo-striate paths described by Du et al. (Du, Li et al. 2023), we performed three iterations of CTT with the SUB, mPFC, or LS as obligatory waypoints: we seeded streamlines from CA1 and retained only those that passed through that waypoint, excluding streamlines that contacted the other two regions. All three CA1-striatum routes exhibited a striosome-favoring bias. In the subiculum pathway, the CA1 volume biased toward striosome-like voxels was 361 mm^3^, whereas CA1 volume biased toward matrix-like voxels was 142 mm^3^ (p<10^-100^; striosome/matrix ratio = 2.55). The medial prefrontal cortex route also exhibited a striosome-dominant pattern, with 321 mm^3^ biased towards striosome-like and 171 mm^3^ biased towards matrix-like voxels (p<10^-100^; striosome/matrix ratio = 1.88). The lateral septum route also demonstrated higher striosome-like bias, with 359 mm^3^ routed towards striosome-like and 143 mm^3^ routed towards matrix-like voxels (p<10^-100^; striosome/matrix ratio = 2.52).

### Striatal Voxels with High Connectivity to CA1 are Striosome-like

As a control, we performed classification targets tractography from striatum to CA1 to confirm our previous results (B-to-A, to compare with our previous A-to-B findings). In each subject we divided the striatum into high-connectivity (uppermost 25% of voxels, those above the 75th percentile of streamline counts) and low-connectivity (lowest 75%) voxels, then extracted the striosome-like bias within those high- and low-connectivity masks (Figure 2). Striatal voxels with the highest amplitude of streamlines that reached CA1 were found to be markedly enriched in striosome-like bias. For the SUB route, the mean striosome bias in high-connectivity voxels was double the bias in low-connectivity voxels (0.68 [SEM: 0.027] vs. 0.32 [SEM: 0.018], respectively; p<10^-100^). For the mPFC route, the mean striosome-bias was 45% higher in high-connectivity than in low-connectivity voxels (0.48 [SEM: 0.038] vs. 0.33 [SEM: 0.018], respectively; p<10^-100^).

**Figure 2:**
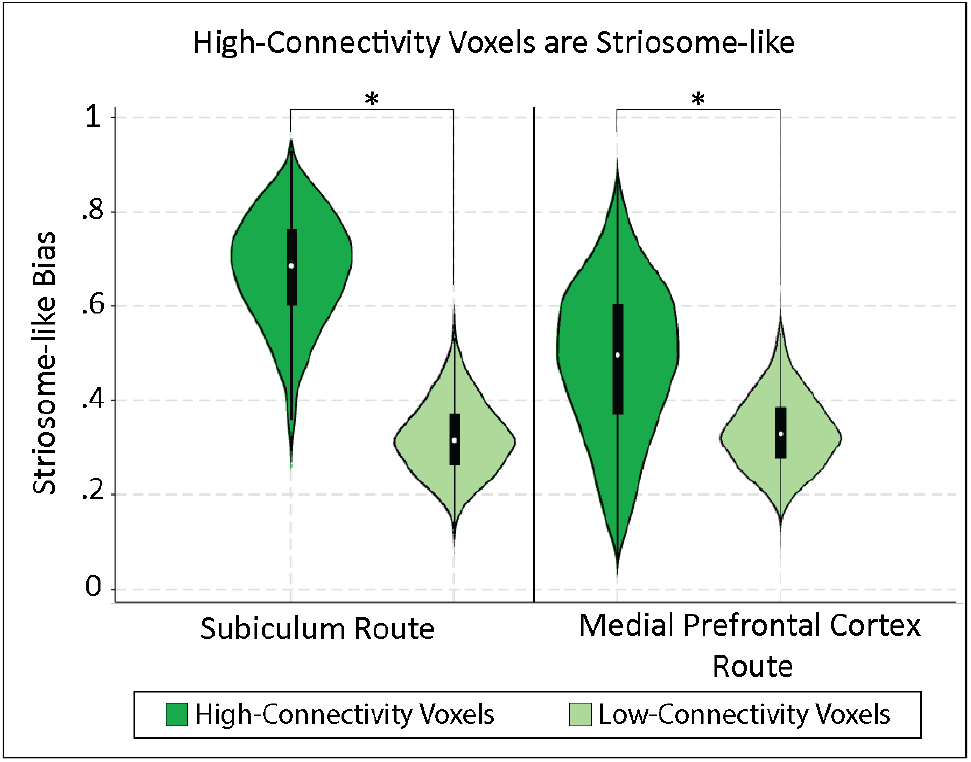
Striatal voxels with high connectivity to CA1 are more striosome-biased than voxels with low connectivity to CA1 for both the subiculum and medial prefrontal cortex routes. Black rectangles illustrate the interquartile range, white points in each rectangle indicate the median, and kernel density plots illustrate the distribution of bias within the cohort. *, p<10^-100^.

### Distinct Connectivity Patterns across Hippocampal Routes and CA1 Subregions

To assess if the SUB and mPFC routes reached distinct parts of the striatum, we assessed the DSC for high-connectivity striatal voxels (uppermost 25^th^ percentile of voxels by streamline count) and low-connectivity striatal voxels (CTT with striatum as seed, CA1 as target) for the SUB and mPFC routes (high, SUB vs. mPFC; low, SUB vs. mPFC). The DSC for high-connectivity voxels was low, suggesting that these CA1 projections reached largely distinct zones within the striatum. The DSC between the SUB and mPFC routes was 17% (p<1×10^-100^, SEM: 0.36%) for high-connectivity voxels and 73% (p<1×10^-100^, SEM: 0.26%) for low-connectivity voxels (Figure 3).

**Figure 3:**
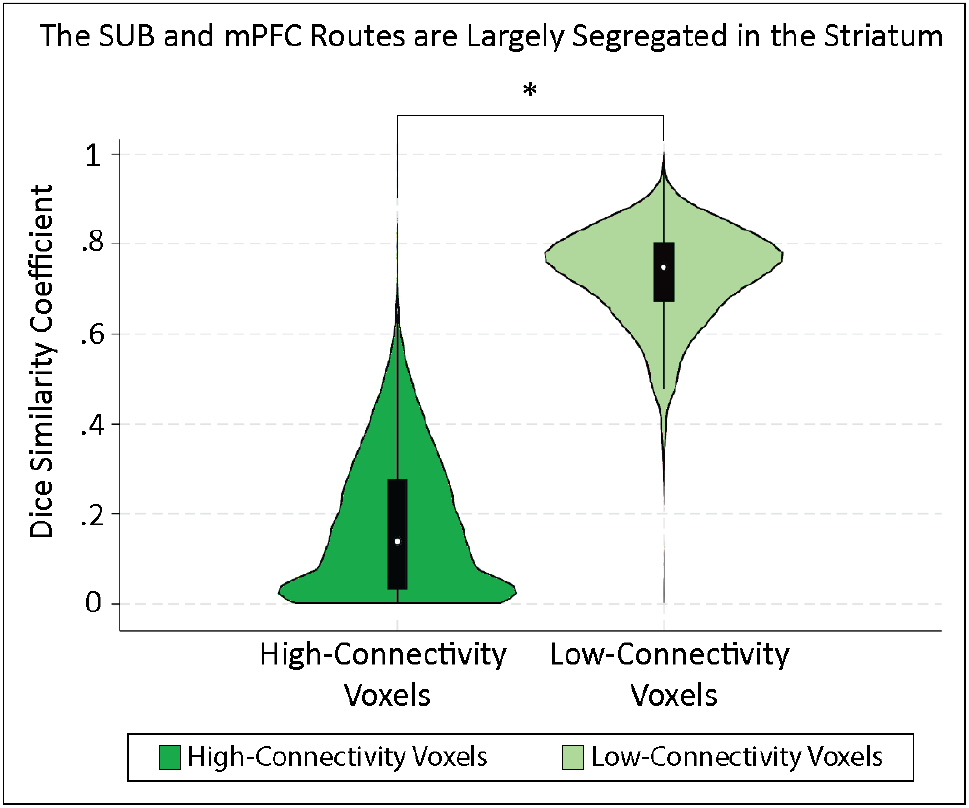
CA1-to-dorsal striatum projections that routed through the subiculum (SUB) were largely segregated, in their striatal terminations, from projections that routed through the medial prefrontal cortex (mPFC). The low DSC in high-connectivity voxels (17.2%, SUB vs. mPFC) and high DSC in low-connectivity voxels (72.9%) indicate that high-precision streamlines in the SUB and mPFC routes terminate in different parts of the striatum. Black rectangles illustrate the interquartile range, white points in each rectangle indicate the median, and kernel density plots illustrate the distribution of bias within the cohort. *, p<1×10^-100^.

To determine if the pathways to the matrix-favoring and striosome-favoring regions of CA1 were distinct, we extracted the DSC for projections between high-connectivity voxels and the matrix-favoring vs. striosome-favoring regions of CA1 (Figure 1). For the SUB route, the DSC between voxels with high connectivity projections to the matrix-favoring vs. striosome-favoring parts of CA1 was 25% (SEM: 0.51%). For the mPFC route, this DSC was 5.9% (SEM: 0.26%). The low DSCs indicate that for each route, the parts of striatum reached by matrix-favoring CA1 were spatially segregated from the parts of striatum reached by striosome-favoring CA1.

### Biases in Functional Connectivity: Correlation Between CA1 and Compartment-like Voxels

Next, we asked whether the structural connectivity networks identified through diffusion MRI predicted functional coupling between CA1 and the striatal compartments. We assessed the resting state functional connectivity between CA1 and the compartment-like masks. In both hemispheres, functional activation in CA1 was strongly correlated with activation in striosome-like voxels and anti-correlated with activation in matrix-like voxels (Figure 4). Right CA1 demonstrated robust positive correlation with striosome-like voxels and negative correlation with matrix-like voxels (striosome: 0.04, matrix = -0.038; CA1-striosome vs. CA1-matrix, p=2.6x10^-12^), while left CA1 showed a similar but weaker bias (striosome: 0.02, matrix: -0.02; CA1-striosome vs. CA1-matrix, p=2.8x10^-4^). These findings suggest a hemispheric asymmetry, with CA1-compartment functional correlations on the right doubling those on the left.

**Figure 4:**
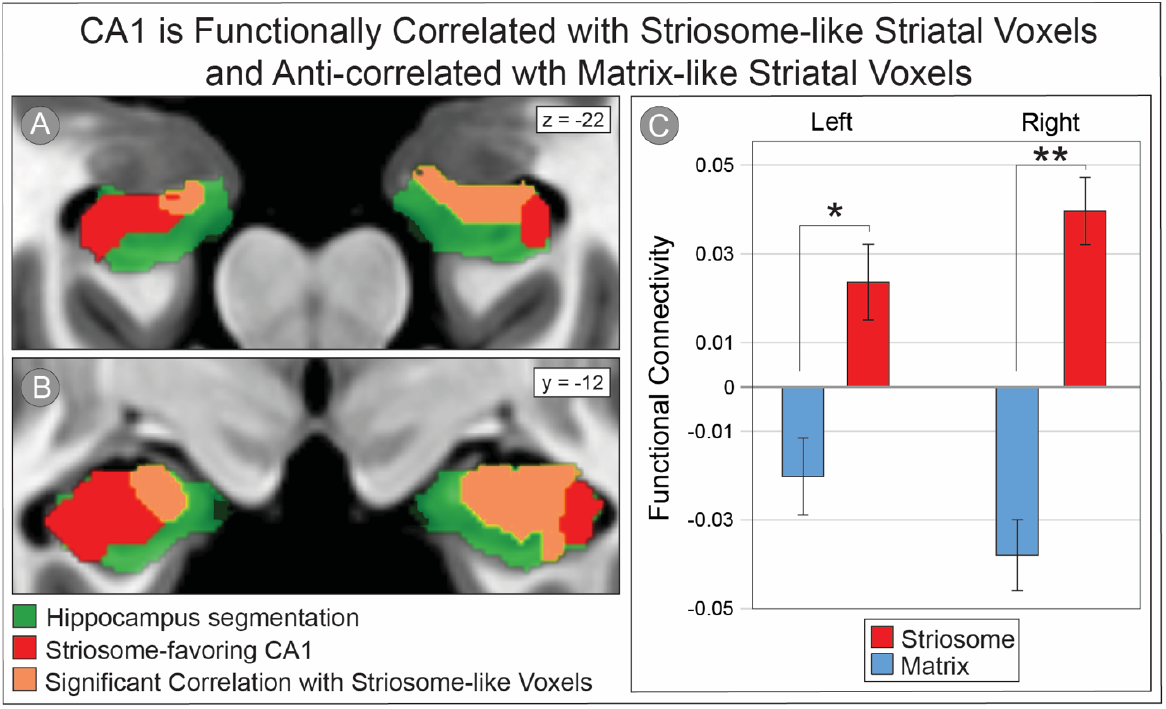
Striosome- and matrix-like masks, defined through structural connectivity, had opposing functional connectivity with hippocampal CA1. In axial (A) and coronal (B) sections, the striosome-favoring zone within CA1 (red) is overlaid on the hippocampus (green voxels). CA1 voxels with significant positive correlation with striosome-like voxels (orange, with rim of yellow to improve visual contrast), which fell within the striosome-favoring zone of CA1. Mean functional connectivity with CA1 (C) are shown for striosome-like (red bars) or matrix-like voxels (blue bars), plotted separately for the left and right hemispheres. Functional activation in striosome-like voxels positively correlated CA1, whereas activation in matrix-like voxels negatively correlated with CA1, revealing a robust compartment-specific dissociation in hippocampal–striatal functional connectivity. Error bars indicate standard error of the mean. The images adhere to radiographic convention and coordinates follow MNI convention. *, p<1×10^-3^; **, p<1×10^-11^.

### Compartment Specificity in CA1-Nucleus Accumbens Connectivity Diverges by Hemisphere

We used CTT to evaluate compartment bias in projections from CA1 to matrix-like and striosome-like voxels in the nucleus accumbens (NAcc). In a combined-hemispheres analysis, the mean volume of CA1 that routed to matrix-like and striosome-like voxels in NAcc was 169 mm^3^ and 134 mm^3^, respectively (p=2.4x10^-10^). The indeterminate volume (0.45< P <0.55) was 29.4 mm^3^. We then assessed voxelwise differences in compartment-like bias in CA1-accumbens structural connectivity (Figure 5). As a control, we segmented the region which contained the main streamline bundle and included this region as an exclusion mask for streamline generation. Removal of the dominant CA1-nucleus accumbens streamline bundle eliminated the previously observed bilateral compartment effect, indicating that this effect was primarily driven by that pathway. In this negative control experiment, CTT now revealed that CA1 connectivity in both the left (striosome: 23.4 mm^3^ vs. matrix: 6.4 mm^3^; p=9.2×10^-54^) and right (striosome: 20.5 mm^3^ vs. matrix: 13.5 mm^3^; p=1.6×10^-8^) hemispheres were now striosome biased.

**Figure 5:**
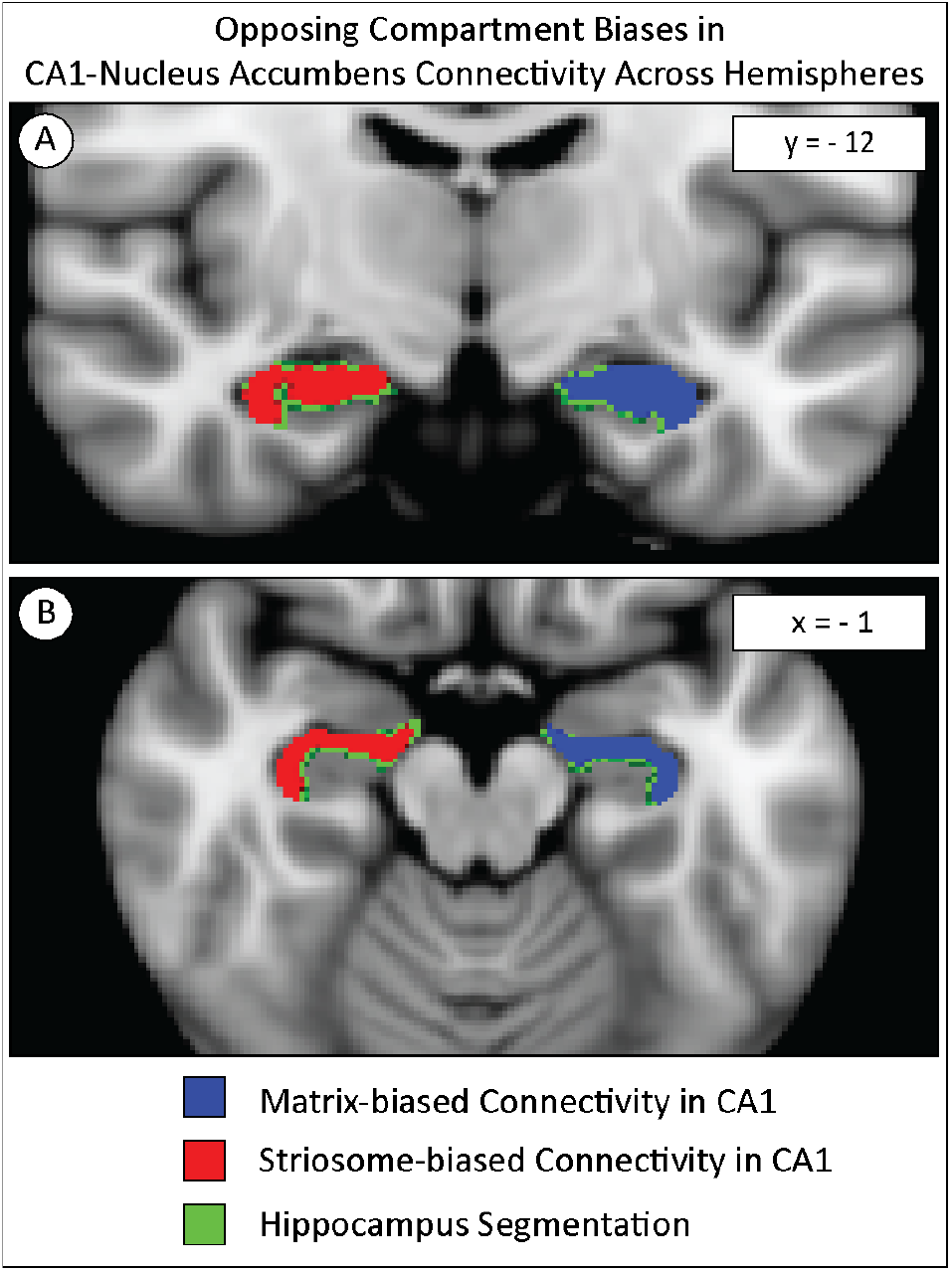
CA1-Nucleus accumbens structural connectivity exhibited compartment biases that were opposite in left and right hemispheres. Streamlines seeded from the right CA1 preferentially contacted striosome-like voxels, with 78% of CA1 voxels showing a striosome bias, whereas streamlines seeded from the left CA1 preferentially contacted matrix-like voxels, with 68% of CA1 voxels showing a matrix bias (coronal, A; axial, B). Significance threshold, p< 0.05, FWE-corrected within randomise. The images adhere to radiographic convention and coordinates follow MNI convention.

CA1 voxels that were biased toward striosome-like masks were 3.1-fold more abundant than those biased toward matrix-like voxels (striosome: 386 mm^3^ vs. matrix: 125 mm^3^; p<10^-100^). Both the left (striosome: 372 mm^3^, matrix: 125 mm^3^; p<10^-100^) and right (striosome: 398 mm^3^, matrix: 125 mm^3^; p<10^-100^) hemispheres exhibited the same pattern of increased striosome-like bias. Voxelwise assessment of bias (using FSL’s *randomise*, Figure 1) identified a symmetrical segment of caudo-lateral CA1 that was significantly biased toward striosome-like voxels (mean probability of connection to striosome-like masks: 0.70), whereas a smaller rostro-medial cluster was biased toward matrix-like voxels (mean probability of connection to matrix-like masks: 0.59; striosome-favoring vs. matrix-favoring volume, p=7.9×10^-92^).

### Route-Specific CA1 to Compartment-like Voxels in the Nucleus Accumbens

To further delineate the polysynaptic pathways to the nucleus accumbens found by Du et al. (Du, Li et al. 2023), we selected streamlines that contacted either the subiculum, lateral septum, or medial prefrontal cortex as an obligatory waypoint, and excluded streamlines that contacted either of the other two regions. This approach was a direct parallel for that utilized above for the dorsal striatum. We extracted the volume of CA1 that was biased to either compartment (P>0.55) for each route (Table 2). Across all three routes, left hemisphere tractography was biased towards matrix-like voxels while the right hemisphere was biased towards striosome-like voxels.

**Table 2:** CA1 to nucleus accumbens tractography exhibits lateralized compartmental biases; the left hemisphere is matrix biased while the right hemisphere is striosome biased.

| Route | Hemisphere | CA1 to Matrix-like Volume | CA1 to Striosome-like Volume | Striosome/Matrix Ratio |
| --- | --- | --- | --- | --- |
| Medial Prefrontal Cortex | Left | 108.3 mm<sup>3</sup> | 44.7 mm<sup>3</sup> | 0.41 |
|  | Right | 65.2 mm<sup>3</sup> | 105.6 mm<sup>3</sup> | 1.6 |
| Subiculum | Left | 159.5 mm<sup>3</sup> | 72.9 mm<sup>3</sup> | 0.46 |
|  | Right | 88.4 mm<sup>3</sup> | 139.8 mm<sup>3</sup> | 1.6 |
| Lateral Septum | Left | 196.4 mm<sup>3</sup> | 64.9 mm<sup>3</sup> | 0.33 |
|  | Right | 95.9 mm<sup>3</sup> | 139.9 mm<sup>3</sup> | 1.5 |

## Discussion

The CA1 subfield is anatomically and functionally segregated along the dorsoventral axis in mammals (Dong, Swanson et al. 2009, Strange, Witter et al. 2014, Tao, Wang et al. 2021). In rodents, dorsal CA1 is preferentially involved in spatial and cognitive processing (Rogers and Kesner 2006, Kim, Brager et al. 2018), whereas ventral CA1 is more closely associated with emotion and behavioral learning (Ruediger, Spirig et al. 2012, McDonald, Balog et al. 2018). This dorsoventral functional gradient is supported by regionally segregated biases in connectivity and spatially patterned gene expression along the hippocampal axis (Dong, Swanson et al. 2009, Fanselow and Dong 2010). Although the human hippocampus is oriented rostro-caudally, rather than dorsoventrally as in the rodent, our findings parallel these dorsoventral organizational schema identified in rodents: lateral-dorsal CA1 showed a bias toward striosome-like voxels, whereas medial-rostral CA1 preferentially targeted matrix-like voxels. The spatial segregation of these projections, both within CA1 and in the striatum, suggests that compartment-specific hippocampal projections in humans may also serve distinct functions. To the best of our knowledge, this type of hippocampal-striatal specialization has not been described previously.

The dorsoventral projection biases we identified suggest a link between regional hippocampal specialization and compartment-specific CA1-striate circuits. Consistent with this, the striosome compartment is selectively vulnerable in Huntington’s disease, and its degeneration has been correlated with the severity of mood dysfunction (Tippett, Waldvogel et al. 2007). Moreover, selective stimulation of the striosome in primates induces pessimistic decision-making (Amemori, Amemori et al. 2020), and striosome-specific cortico-striatal circuits have been shown to modulate cognitive and emotion-related functions (Friedman, Homma et al. 2015). In rodents, ventral CA1 has been strongly implicated in mood regulation and learning (Ruediger, Spirig et al. 2012, McDonald, Balog et al. 2018). When accounting for the rotation of hippocampal axes between species, our findings suggest that the spatial equivalent of the rodent ventral CA1 (dorsolateral CA1 in humans) may preferentially engage striosome-specific circuitry, providing an anatomical basis for mood modulation by CA1-striosome specific circuitry. By contrast, in humans rostral-medial CA1, the spatial equivalent of the rodent dorsal CA1, showed preferential connectivity with the matrix compartment, where increased connectivity with sensorimotor zones may underlie a functional specialization in visual and spatial processing (Flaherty and Graybiel 1993, Flaherty and Graybiel 1994).

Interactions between the intermediate regions (mPFC, SUB, or LS) with hippocampal CA1 are important for memory functions. For example, lesioning the mPFC and disrupting the mPFC-CA1 circuit decreased memory formation in rats (Schlichting and Preston 2014, Eichenbaum 2017). The SUB is a major output target of the hippocampal formation (Xu, Sun et al. 2016) and is a central region which acts as a recipient, comparator, and distributor of processed information within the hippocampal memory system (Naber, Witter et al. 2000). The LS is hypothesized to process the spatial information from CA1 and forward this information to downstream targets for the purposes of reinforcement and reward seeking behavior (Wirtshafter and Wilson 2020). We found in humans that all three routes were striosome-biased, which may reflect the striosome’s role in modulating cognition (Beste, Moll et al. 2018).

The opposing compartmental biases observed in left and right CA1 projections to the nucleus accumbens may reflect underlying hemispheric asymmetries in hippocampus-nucleus accumbens connectivity. Supporting this possibility, probabilistic tractography in adolescents has shown that hippocampal connectivity to the nucleus accumbens is approximately twice as strong in the left hemisphere as in the right (Zabri SH, Ahmad AH et al. 2023). Given the association of the striosome with mood regulation (Tippett, Waldvogel et al. 2007), such lateralized circuitry may also align with hypotheses of right-hemisphere dominance in emotional processing (Gainotti 2019, Gainotti 2019). Histochemical and transcriptomic studies support compartmental organization within the nucleus accumbens (Jongen-Rĕlo, Groenewegen et al. 1993, Tanimura, King et al. 2011, Reiner, Chehimi et al. 2024, Contesse, Bektash et al. 2025), but compartment-specific connectivity biases in this region remain largely unexplored. Our results therefore provide a first suggestion that compartmental biases within the nucleus accumbens are lateralized, a finding that requires replication and further exploration in animal models.

Our findings have several important limitations. First, connectivity-based striatal parcellation relies on probabilistic tractography, an inferential technique that cannot distinguish afferent from efferent projections, is susceptible to both false positives and false negatives, and is substantially influenced by acquisition parameters, all of which may affect estimates of compartment-like bias (Jones and Cercignani 2010). Our cohort is derived from the HCP S1200, whose scanner protocols were standardized to minimize the effect of scanner variations. Second, our selection of bait regions depends on tract-tracer data from animals for compartment-specific connectivity. Because neural circuitry may differ between species, these comparisons should be interpreted with caution. Third, the spatial resolution of our diffusion voxels matched the maximum diameter of striosome branches (∼1.25 mm; (Graybiel and Ragsdale 1978, Holt, Graybiel et al. 1997)), meaning that even voxels with high striosome-like bias contained some matrix tissue. Higher resolution diffusion MRI could provide greater granularity, potentially isolating voxels composed purely of matrix or striosome tissue. Despite these limitations, our method demonstrated a low test–retest error (0.14%; (Waugh, Hassan et al. 2022)), and the relative abundance and spatial distribution of compartment-like voxels correspond closely with histological descriptions of matrix and striosome (Funk, Hassan et al. 2023, Funk, Hassan et al. 2024, Tieu and Waugh 2025, Waugh, Hassan et al. 2025). Nonetheless, it is important to emphasize that this method is inferential and cannot substitute for direct identification of the compartments in striatal tissue. Although our analyses sought to further delineate the hippocampus-nucleus accumbens pathways reported by Du et al. (Du, Li et al. 2023), it should be noted that our compartmental-favoring bait regions were identified based on structural connectivity with the dorsal striatum. The optimal bait regions for identifying matrix and striosome in the nucleus accumbens may differ, but we could identify no prior animal studies that mapped compartment-specific projections with the nucleus accumbens.

In conclusion, this study delineates the polysynaptic circuitry from hippocampal CA1 to the striatal compartments in humans by identifying biases in structural connectivity connections. Dissecting the architecture of these circuits is critical for understanding how information is transmitted and transformed in the process of memory encoding, recall, and learning. Among these networks, the hippocampus and striatum interact extensively to support memory and behavioral learning (Mizumori, Yeshenko et al. 2004); Rossato, 2006 #3293; Müller, 2018 #3298}. Selective projections to the striatal compartments have the potential to fundamentally shape how information is processed during learning and reward. Our findings provide evidence that projections from CA1 to the striatum are compartmentally organized, exhibiting spatially segregated biases toward the matrix and striosome compartments.

## Data availability statement

Publicly available datasets were analyzed in this study. HCP data can be found here: https://www.humanconnectome.org/study/hcp-young-adult/document/1200-subjects-data-release. This dataset is BIDS compliant. The code, bait, seed, and exclusion masks necessary to complete striatal parcellation can be accessed here: github.com/jeff-waugh/Striatal-Connectivity-based-Parcellation.

## Author Contributions

AT: data acquisition and analysis, initial manuscript drafting, and critical revision of the manuscript.

AS: data analysis, critical revision of the manuscript

JW: data acquisition, analysis, and interpretation, initial manuscript drafting, and critical manuscript revision.

All authors contributed to the article and approved the final version for submission.

